# Using high-resolution functional MRI to differentiate impacts of strabismic and anisometropic amblyopia on evoked ocular dominance activity in humans

**DOI:** 10.1101/2024.02.11.579855

**Authors:** Shahin Nasr, Jan Skerswetat, Eric D. Gaier, Sarala N. Malladi, Bryan Kennedy, Roger B.H. Tootell, Peter Bex, David G. Hunter

**Affiliations:** Athinoula A. Martinos Center for Biomedical Imaging, Massachusetts General Hospital, Charlestown, MA, United States; Department of Radiology, Harvard Medical School, Boston, MA, United States; Department of Psychology, Northeastern University, Boston, MA, United States; Department of Ophthalmology, Harvard Medical School, Boston, MA, United States; Department of Ophthalmology, Boston’s Children Hospital, Boston, MA, United States; Picower Institute for Learning and Memory, Massachusetts Institute of Technology, Cambridge, MA, United States

**Keywords:** Amblyopia, Monocular Response, Interocular Visual Acuity Difference, Columnar Organization, High-Resolution FMRI

## Abstract

We employed high-resolution functional MRI (fMRI) to distinguish the impacts of anisometropia and strabismus (the two most frequent causes of amblyopia) on the evoked ocular dominance (OD) response. Sixteen amblyopic participants (8 females), comprising 8 individuals with strabismus, 7 with anisometropia, 1 with deprivational amblyopia, along with 8 individuals with normal visual acuity (1 female), participated in this study for whom, we measured the difference between the response to stimulation of the two eyes, across early visual areas (V1-V4).

In controls, as expected from the organization of OD columns, the evoked OD response formed a striped pattern that was mostly confined to V1. Compared to controls, the OD response in amblyopic participants formed larger fused patches that extended into downstream visual areas. Moreover, both anisometropic and strabismic participants showed stronger OD responses in V1, as well as in downstream visual areas V2-V4. Although this increase was most pronounced in V1, the correlation between the OD response level and the interocular visual acuity difference (measured behaviorally) was stronger in higher-level visual areas (V2–V4).

Beyond these common effects, and despite similar densities of amblyopia between the anisometropic and strabismic participants, we found a greater increase in the size of V1 portion that responded preferentially to fellow eye stimulation in anisometropic compared to strabismic individuals. We also found a greater difference between the amplitudes of the response to binocular stimulation, in those regions that responded preferentially to the fellow vs. amblyopic eye, in anisometropic compared to strabismic subjects. In contrast, strabismic subjects demonstrated increased correlation between the OD responses evoked within V1 superficial and deep cortical depths, whereas anisometropic subjects did not.

These results provide some of the first direct functional evidence for distinct impacts of strabismus and anisometropia on the mesoscale functional organization of the human visual system, thus extending what was inferred previously about amblyopia from animal models.

## 1. Introduction

Ocular dominance (OD), the preference for responding to stimulation of one eye over the other, is a prominent characteristic of most neurons in primary visual cortex (V1) (Hubel and Wiesel, 1962). In humans and many non-human mammals, neurons with similar OD preferences are grouped together in ocular dominance columns (ODCs), which form a fundamental architectural feature of V1 (LeVay et al., 1975; Tootell et al., 1988; Sincich et al., 2003; Adams et al., 2007). The development of ODCs depends on balanced binocular visual input at early life stages, also known as the critical period (Hubel et al., 1977; LeVay et al., 1980; Horton and Hocking, 1997). Perturbations to normal binocular visual experience during the critical period impact the selectivity and distribution of ODCs and is associated with amblyopia, a prevalent neurodevelopmental disorder affecting a range of visual functions in one or both eyes (McKee et al., 2003; Maurer and McKee, 2018).

Much of our current understanding of amblyopia and its impact on ODCs is based on electrophysiological and anatomical studies conducted in animal models (Fig. 1). According to these studies, asymmetric binocular vision in early life stages, caused either by misalignment of the eyes (strabismus), differential optics of the eyes (anisometropia), or monocular deprivation, leads to a reduction in the number of V1 neurons that respond binocularly (Crawford and Von Noorden, 1979; Crawford et al., 1996; Smith III et al., 1997b; Kiorpes et al., 1998; Bi et al., 2011). Beyond this common effect, anisometropia and strabismus may impact the evoked OD response in different ways. Specifically, anisometropia, even in milder forms, is associated with a decrease in the number of neurons that respond preferentially to the amblyopic eye. Whereas such a bias is only detectable in strabismic participants with severe amblyopia (Crawford et al., 1996; Kiorpes et al., 1998; Bi et al., 2011). Moreover, strabismus (but not anisometropia) increases the segregation between ODCs with opposing ocular preference (Lowel, 1994; Tychsen et al., 2004).

**Figure 1.**
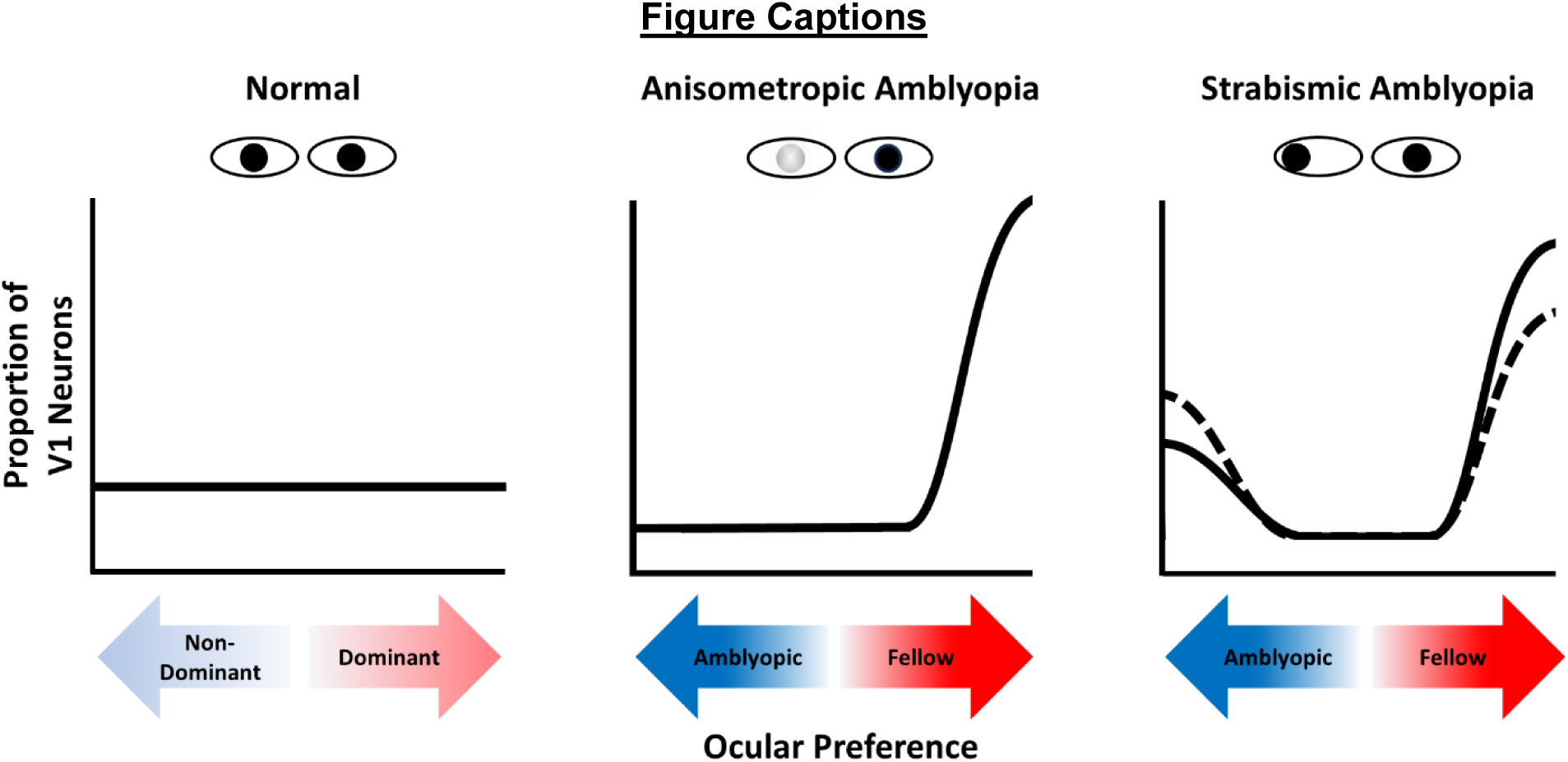
Schematic representation of the relative impact of anisometropic and strabismic amblyopia on the ocular preference of V1 neurons in non-human primates. Individuals with normal binocular vision and no amblyopia (**left**) have a uniform preference for either eye, with some neurons favoring the dominant or non-dominant eye and others showing varying degrees of binocular preference. Amblyopic individuals (regardless of cause) show a decrease in the total number of binocular neurons in V1 (Crawford and Von Noorden, 1979; Crawford et al., 1996; Smith III et al., 1997b; Kiorpes et al., 1998; Bi et al., 2011), while the distribution varies with type: In anisometropic amblyopia (**middle**), this effect is accompanied with a decrease in the number of V1 neurons that respond preferentially to the amblyopic eye, even in those with milder forms of amblyopia. In strabismic individuals with milder forms of amblyopia (**right - solid line**), amblyopic eye-preferring neurons remain frequently detectable across V1, whereas in more severe forms (**right - dashed line**), these neurons are less frequently observed.

In humans, fMRI has been used successfully to localize OD bands within primary visual cortex non-invasively (Menon et al., 1997; Cheng et al., 2001; Yacoub et al., 2007; Nasr et al., 2016). Using this technique, further studies suggest amblyopia is associated with a greater number of voxels responding preferentially to the fellow eye compared to the amblyopic eye (Algaze et al., 2002; Goodyear et al., 2002; Liu et al., 2004), and that the OD activity was stronger in amblyopic participants compared to controls (Conner et al., 2007). It was also suggested that amblyopia changes the mechanism of binocular interaction from excitation to suppression (Farivar et al., 2011; Thompson et al., 2019). However, these studies did not clarify whether this effect extends throughout all of V1 or if this effect is limited to regions that response preferentially to the amblyopic eye. Further, these studies did not distinguish the impacts of anisometropia vs. strabismus on the evoked OD response and/or the mesoscale functional organization of V1, presumably due to limited spatial resolution and contrast-to-noise ratio of the neuroimaging techniques available at the time.

To address these knowledge gaps, this study used higher spatial resolution fMRI (voxel size = 1 mm isotropic), conducted in a 7T MR scanner. Advanced neuroimaging technologies were used to mitigate the contribution of different nuisance factors (e.g., cardiac and respiratory activities) on signal quality (Polimeni et al., 2015). Additionally, the contrast-to-noise ratio was improved by minimizing the level of unwanted signal blurring without applying any spatial smoothing within cortical layers (Blazejewska et al., 2019; Wang et al., 2022).

Using these methods, first we compared the impact(s) of strabismus and anisometropia on the spatial distribution and columnar organization of the evoked OD response in human V1 Notably, while it is known that amblyopia changes the selectivity level of ‘horizontal’ (i.e., surface-parallel) connection between ODCs (Tychsen et al., 2004), the impact of amblyopia on ‘radial’ (i.e., perpendicular to the surface) connections between cortical layers remains mostly unknown even in animal models (Horton and Hocking, 1997). Second, we measured the impact of amblyopia on the amplitude of OD responses in V1, and in downstream extrastriate visual areas (V2-V4). Third, we aimed to compare the correlation between the evoked OD response and the interocular visual acuity difference as a measure of amblyopia severity across the human visual system hierarchy. Lastly, we aimed to compare the evoked activity across V1 regions to binocular stimulation to test whether the binocular response varies between regions that respond preferentially to the fellow vs. amblyopic eye. Our findings provided the first direct evidence for the differential impact of anisometropia and strabismus on the OD response across different visual areas and confirmed the hypothesized link between the evoked OD response and the interocular visual acuity difference in amblyopia.

## 2. Results

The OD response was measured in 24 human participants, 16 with amblyopia caused either by strabismus (*n=*8), anisometropia (*n=*7) or deprivational amblyopic (*n=*1), and 8 control participants with normal or corrected-to-normal vision. In addition to data from these individuals, we also measured the OD response in one strabismic (but non-amblyopic) participant whose data are presented separately. To measure the evoked response to stimulation of the eyes, each participant was scanned twice on different days. During these scans, moving random dots were presented to each eye separately (using anaglyphic goggles) in a blocked-design paradigm (see Methods). The OD response was measured for each participant by averaging the activity evoked across these two sessions and calculating the (absolute) difference between the response to stimulation of dominant/fellow vs. non-dominant/amblyopic eye. A subset of subjects (Table 1) also participated in a control test to measure responses to dichoptically presented grating stimuli. Outside the scanner, all participants were tested to measure their visual acuity and stereoacuity, to identify their dominant eye, and to test for suppression and/or diplopia (see Methods).

**Table 1.**
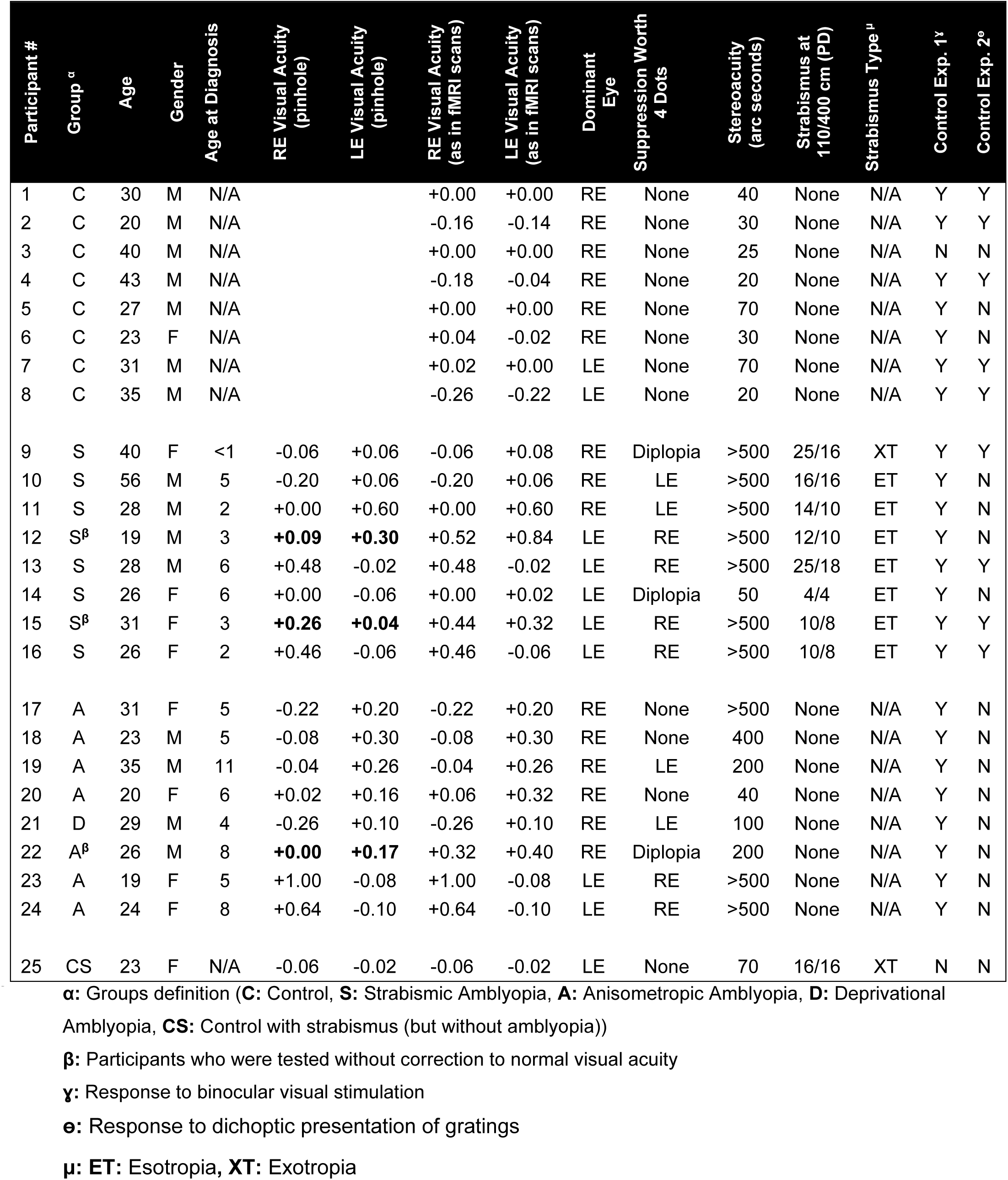
Demography and ophthalmologic assessment of the participants.

| Participant # | Group <sup>α</sup> | Age | Gender | Age at Diagnosis | RE Visual Acuity (pinhole) | LE Visual Acuity (pinhole) | RE Visual Acuity (as in fMRI scans) | LE Visual Acuity (as in fMRI scans) | Dominant Eye | Suppression Worth 4 Dots | Stereoacuity (arc seconds) | Strabismus at 110/400 cm (PD) | Strabismus Type <sup>μ</sup> | Control Exp. 1<sup>γ</sup> | Control Exp. 2<sup>ε</sup> |
| --- | --- | --- | --- | --- | --- | --- | --- | --- | --- | --- | --- | --- | --- | --- | --- |
| 1 | C | 30 | M | N/A |  |  | +0.00 | +0.00 | RE | None | 40 | None | N/A | Y | Y |
| 2 | C | 20 | M | N/A |  |  | -0.16 | -0.14 | RE | None | 30 | None | N/A | Y | Y |
| 3 | C | 40 | M | N/A |  |  | +0.00 | +0.00 | RE | None | 25 | None | N/A | N | N |
| 4 | C | 43 | M | N/A |  |  | -0.18 | -0.04 | RE | None | 20 | None | N/A | Y | Y |
| 5 | C | 27 | M | N/A |  |  | +0.00 | +0.00 | RE | None | 70 | None | N/A | Y | N |
| 6 | C | 23 | F | N/A |  |  | +0.04 | -0.02 | RE | None | 30 | None | N/A | Y | N |
| 7 | C | 31 | M | N/A |  |  | +0.02 | +0.00 | LE | None | 70 | None | N/A | Y | Y |
| 8 | C | 35 | M | N/A |  |  | -0.26 | -0.22 | LE | None | 20 | None | N/A | Y | Y |
| 9 | S | 40 | F | <1 | -0.06 | +0.06 | -0.06 | +0.08 | RE | Diplopia | >500 | 25/16 | XT | Y | Y |
| 10 | S | 56 | M | 5 | -0.20 | +0.06 | -0.20 | +0.06 | RE | LE | >500 | 16/16 | ET | Y | N |
| 11 | S | 28 | M | 2 | +0.00 | +0.60 | +0.00 | +0.60 | RE | LE | >500 | 14/10 | ET | Y | N |
| 12 | S<sup>β</sup> | 19 | M | 3 | **+0.09** | **+0.30** | +0.52 | +0.84 | LE | RE | >500 | 12/10 | ET | Y | N |
| 13 | S | 28 | M | 6 | +0.48 | -0.02 | +0.48 | -0.02 | LE | RE | >500 | 25/18 | ET | Y | Y |
| 14 | S | 26 | F | 6 | +0.00 | -0.06 | +0.00 | +0.02 | LE | Diplopia | 50 | 4/4 | ET | Y | N |
| 15 | S<sup>β</sup> | 31 | F | 3 | **+0.26** | **+0.04** | +0.44 | +0.32 | LE | RE | >500 | 10/8 | ET | Y | Y |
| 16 | S | 26 | F | 2 | +0.46 | -0.06 | +0.46 | -0.06 | LE | RE | >500 | 10/8 | ET | Y | Y |
| 17 | A | 31 | F | 5 | -0.22 | +0.20 | -0.22 | +0.20 | RE | None | >500 | None | N/A | Y | N |
| 18 | A | 23 | M | 5 | -0.08 | +0.30 | -0.08 | +0.30 | RE | None | 400 | None | N/A | Y | N |
| 19 | A | 35 | M | 11 | -0.04 | +0.26 | -0.04 | +0.26 | RE | LE | 200 | None | N/A | Y | N |
| 20 | A | 20 | F | 6 | +0.02 | +0.16 | +0.06 | +0.32 | RE | None | 40 | None | N/A | Y | N |
| 21 | D | 29 | M | 4 | -0.26 | +0.10 | -0.26 | +0.10 | RE | LE | 100 | None | N/A | Y | N |
| 22 | A<sup>β</sup> | 26 | M | 8 | **+0.00** | **+0.17** | +0.32 | +0.40 | RE | Diplopia | 200 | None | N/A | Y | N |
| 23 | A | 19 | F | 5 | +1.00 | -0.08 | +1.00 | -0.08 | LE | RE | >500 | None | N/A | Y | N |
| 24 | A | 24 | F | 8 | +0.64 | -0.10 | +0.64 | -0.10 | LE | RE | >500 | None | N/A | Y | N |
| 25 | CS | 23 | F | N/A | -0.06 | -0.02 | -0.06 | -0.02 | LE | None | 70 | 16/16 | XT | N | N |

**α**: Groups definition (**C**: Control, **S**: Strabismic Amblyopia, **A**: Anisometropic Amblyopia, **D**: Deprivational
Amblyopia, **CS**: Control with strabismus (but without amblyopia))
**β**: Participants who were tested without correction to normal visual acuity
**γ**: Response to binocular visual stimulation
**ε**: Response to dichoptic presentation of gratings
**μ**: **ET**: Esotropia, **XT**: Exotropia

### 2.1. Age and interocular visual acuity difference

Table 1 shows the participant’s demographics and visual testing results. One-way ANOVA (anisometropic vs. strabismic vs. control) did not yield any significant age differences across the three groups (F(2, 23)=1.11, *p*=0.35). As expected, a similar analysis applied to the interocular visual acuity difference showed a significant effect of group (F(2, 23)=8.08, *p*<0.01) driven by the increased interocular visual acuity difference in both anisometropic and strabismic individuals relative to controls (*p*<0.01; Bonferroni corrected for multiple comparison). Interocular visual acuity difference was similar between the anisometropic and strabismic individuals in our participants (*p*=0.89). Visual acuities of amblyopic (*p*=0.29) and fellow (*p*=0.83) eyes were not different between anisometropic and strabismic participants. Thus, age, interocular visual acuity difference, and visual acuity in the amblyopic and fellow eyes were comparable between anisometropic and strabismic individuals.

Monocular suppression and diplopia were more common in strabismic compared to anisometropic participants (Table 1). Also, as expected based on previous studies (Levi et al., 2015), more strabismic individuals demonstrated severely impaired stereoacuity (>500 arc seconds) than anisometropic individuals. All amblyopic individuals had a history of either patching or atropine therapy in childhood.

### 2.2. Head position stability during the fMRI tests

Head motion has a strong impact on the fMRI signal and may influence the level and pattern of evoked fMRI responses which might in turn confound between-group comparisons. Thus, as the first step, we compared the level of head motion between control, strabismic and anisometropic participants. Since all individuals were scanned at least two times on different days, we also tested the consistency of head motion between sessions. One-way repeated measures ANOVA (session (first vs. second)), with a group factor (control vs. strabismic vs. anisometropic individuals), to the measured level of head motion (see Methods) did not yield a significant effect of group (F(2, 21)=0.08, *p*=0.92) or group × session interaction (F(2, 21)=2.57, *p*=0.10) on the degree of head motion. Thus, across the two scan sessions, head motion appears to be comparable across the groups. Head motion was nevertheless included as a nuisance co-variate in all analyses to reduce any residual impact of head motion on our findings.

### 2.3. OD activity mapping

We measured the evoked OD activity for all participants in both deep, middle, and superficial cortical depth levels across visual areas V1-V4 by subtracting the response of the non-dominant eye from the response of the dominant eye. Fig. 2A shows the evoked OD activity in a control participant (Participant #1) across deep, middle and superficial layers. Consistent with post-mortem anatomical studies in humans (Adams et al., 2007) and non-human primates (Hubel et al., 1976; Tootell et al., 1988; Sincich et al., 2003) with normal vision, the cortical topography of the evoked OD response was organized into mostly-parallel stripes. These striped patterns were similarly detected across cortical depths, reflecting the columnar organization of V1 ODCs (Tootell et al., 1988). In both hemispheres, these stripes were predominantly limited to the regions of V1 (r<10°), representing the central retinotopic visual field that were stimulated during the scans. This pattern was consistently observed in all control participants in each hemisphere (Fig. 3).

**Figure 2.**
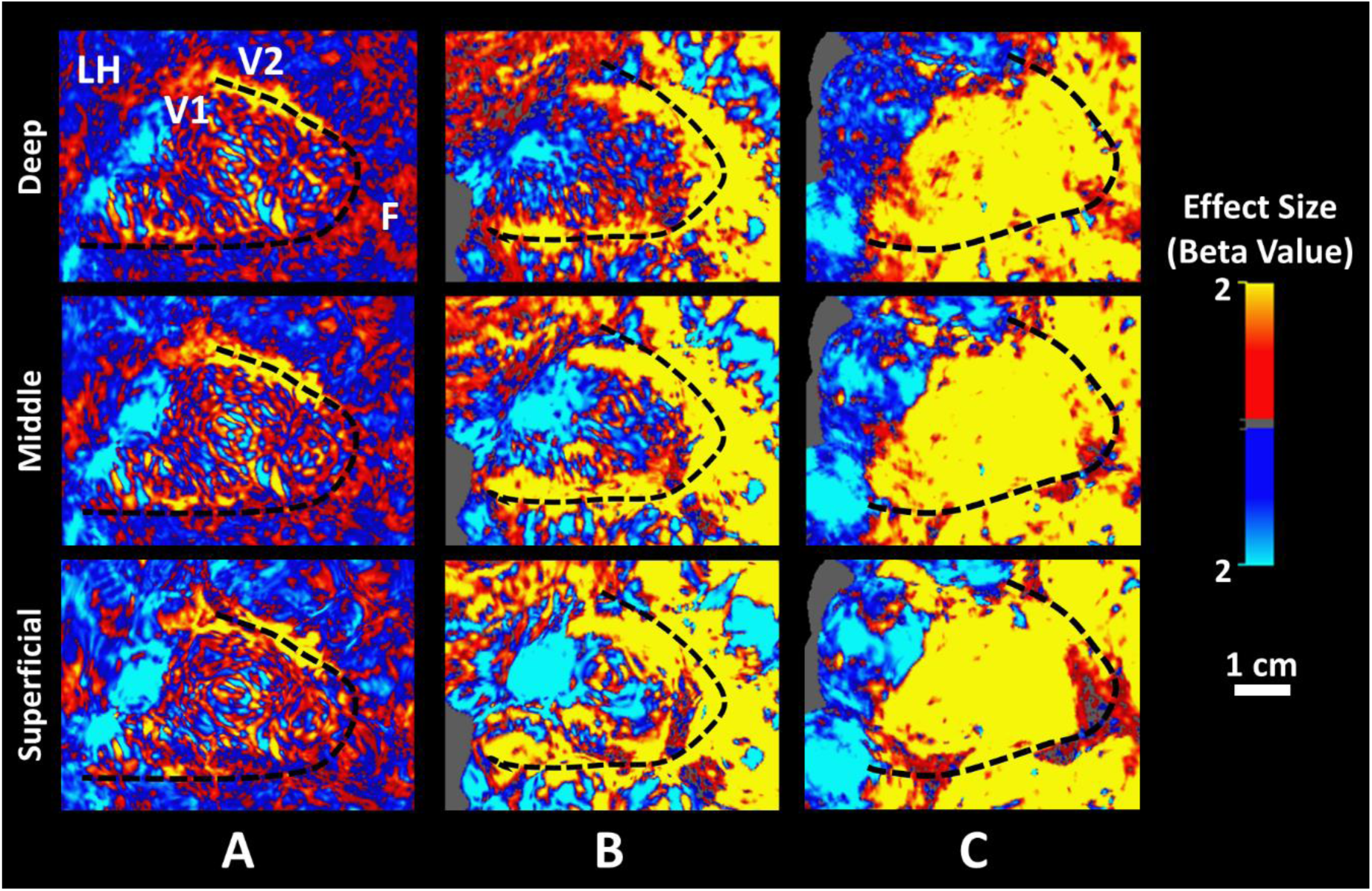
The OD response evoked by contrasting the response, evoked within the left hemisphere (LH), to stimulation of dominant/fellow (red to yellow) vs. non-dominant/amblyopic (blue to cyan) eye across cortical depth levels, measured during two separate scan sessions. Panels **A-C** show the unthresholded activity maps detected within deep (top), middle and superficial (bottom) cortical depths, in the left hemisphere of a control (Subject #1; Table 1), a strabismic (Subject #13), and an anisometropic (Subject #17) subject, respectively. In the control participants, the OD activity formed mostly parallel stripes that were mostly confined to V1 borders. In the amblyopic participants, especially the anisometropic individual, OD stripes were less pronounced, and the evoked activity extended well beyond the V1 border. This phenomenon was apparent comparably detectable across cortical depths. In all panels, activity maps are overlaid on the subject’s own reconstructed cortical surface. The V1-V2 border (black dashed line) is also defined for each subject based on their own retinotopic mapping. The foveal direction is shown with letter F in top-left panel.

**Figure 3.**
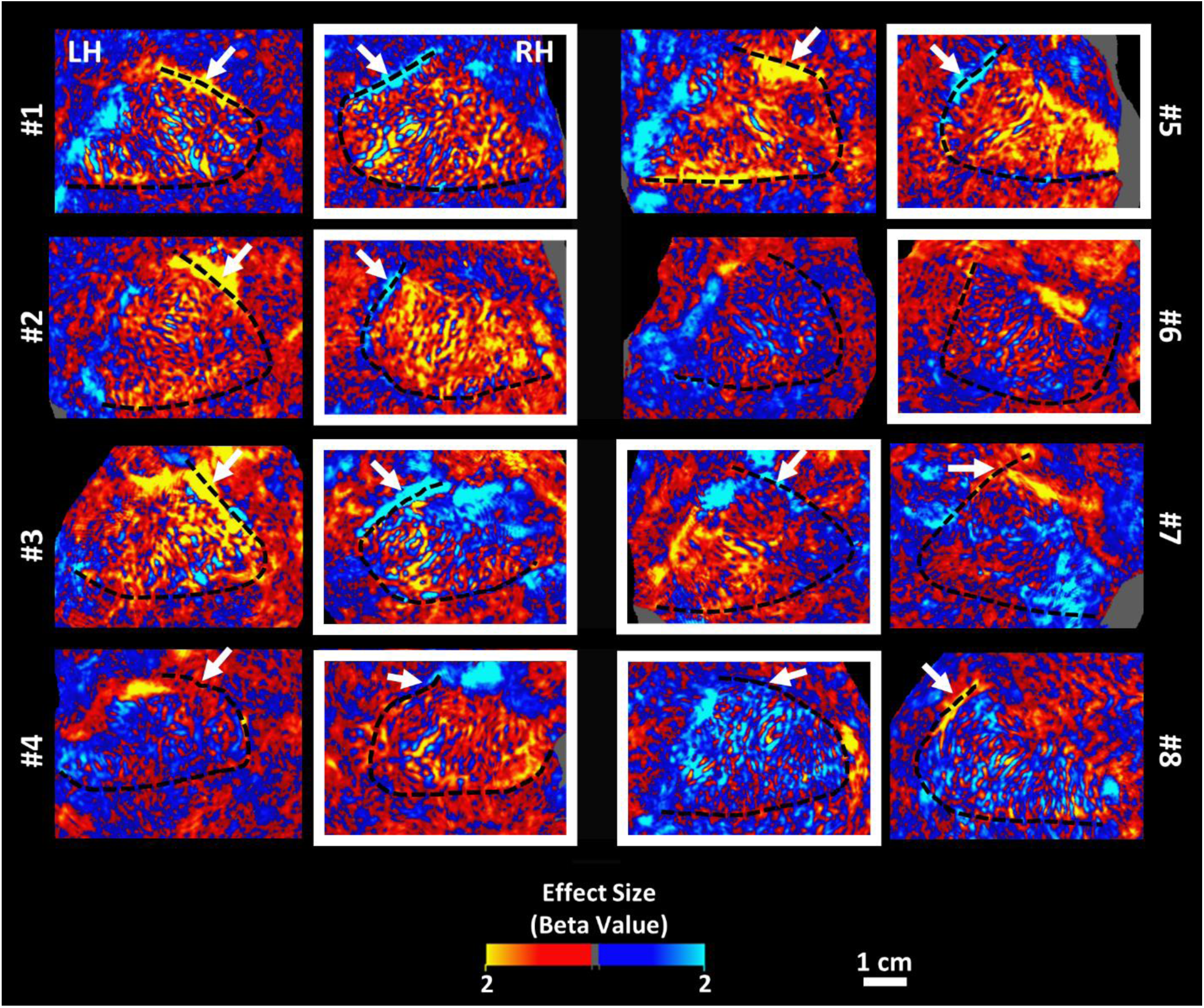
The OD activity mapping in 8 control participants, collected from deep cortical depths. In all participants, the striped pattern was apparent within V1. The response amplitude decreased sharply outside the V1 border. For each subject, the white box indicates the hemisphere ipsilateral relative to the dominant eye. The white arrowheads show the large activity patch along the dorsal portion of V1-V2 that responded preferentially to the contralateral eye. This patch was detectable in almost all participants except for participant #6. Other details are the same as in Fig. 2.

In all controls, we detected a fused activity patch close to the dorsal portion of V1-V2 border that responded preferentially to the contralateral eye (Fig. 3). Notably, this cortical region represents the inferonasal visual field occluded by the head coil resulting in monocular representation by the other eye. Also as expected, we did not detect representation of the blind spot and/or temporal monocular crescent, because these regions are represented more peripherally (r>15°) outside the stimulus borders and scan coverage (Tootell et al., 1998; Awater et al., 2005; Adams et al., 2007; Nasr et al., 2020).

Fig. 2B and 2C illustrate the evoked OD activity in a strabismic (Participant #13; interocular visual acuity differences = 0.50 logMAR) and an anisometropic participant (Participant #17; interocular visual acuity differences = 0.42 logMR), respectively, with comparable levels of interocular visual acuity difference. Compared to controls, OD activity was stronger and formed larger, fused patches at all three cortical depth levels and extended downstream to visual areas V2-V4 (see also Figs. 4 and 5).

**Figure 4.**
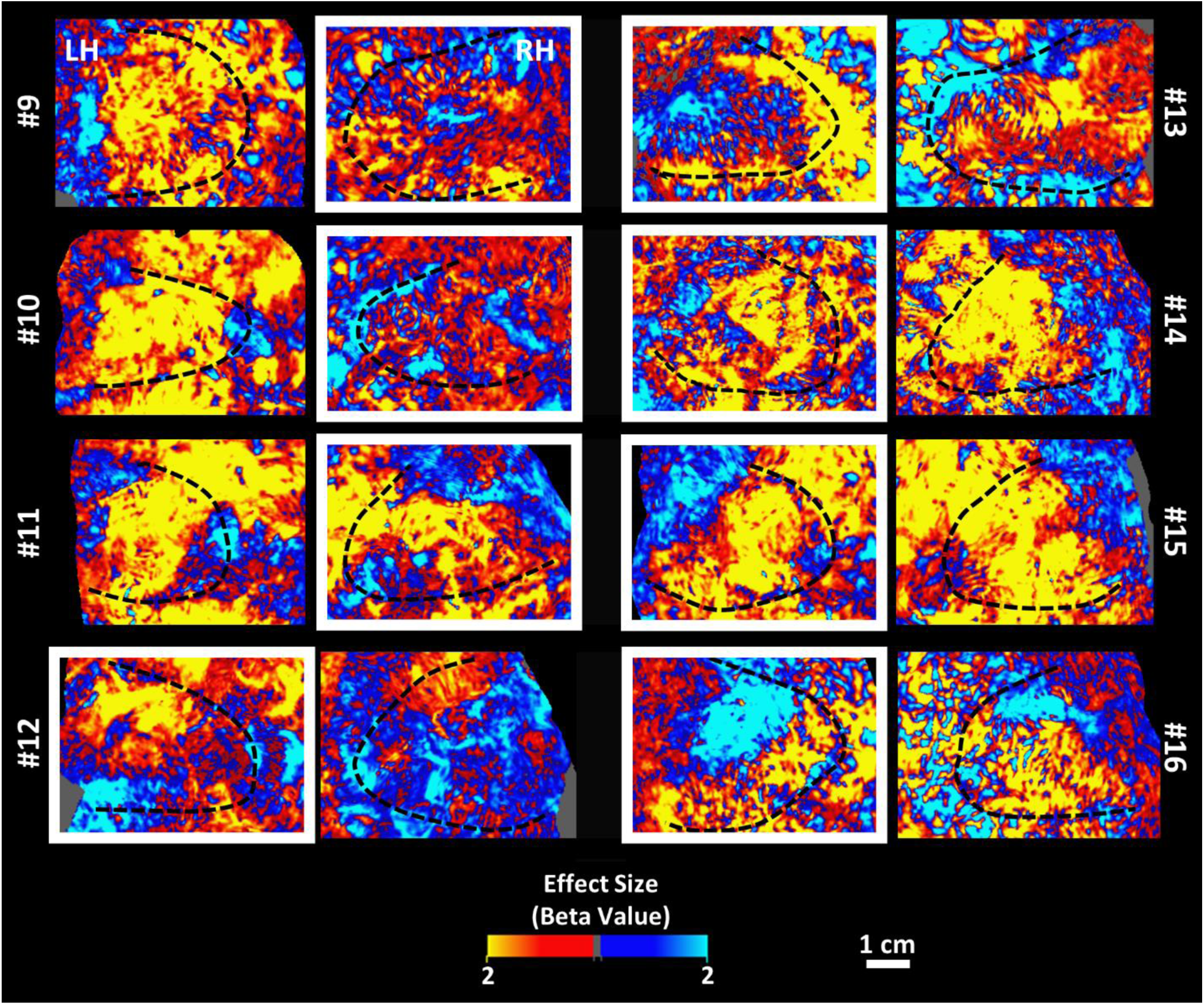
The OD activity mapping in 8 strabismic participants, collected from deep cortical depths. Compared to controls (Fig. 3), the amplitude of OD response was larger. Moreover, the OD response extended beyond the V1-V2 borders into downstream visual areas. This effect was accompanied by an extension of those regions that responded preferentially to the fellow eye. This overrepresentation was more pronounced in the hemisphere contralateral relative to the fellow eye. Other details are the same as in Figs. 2 and 3.

**Figure 5.**
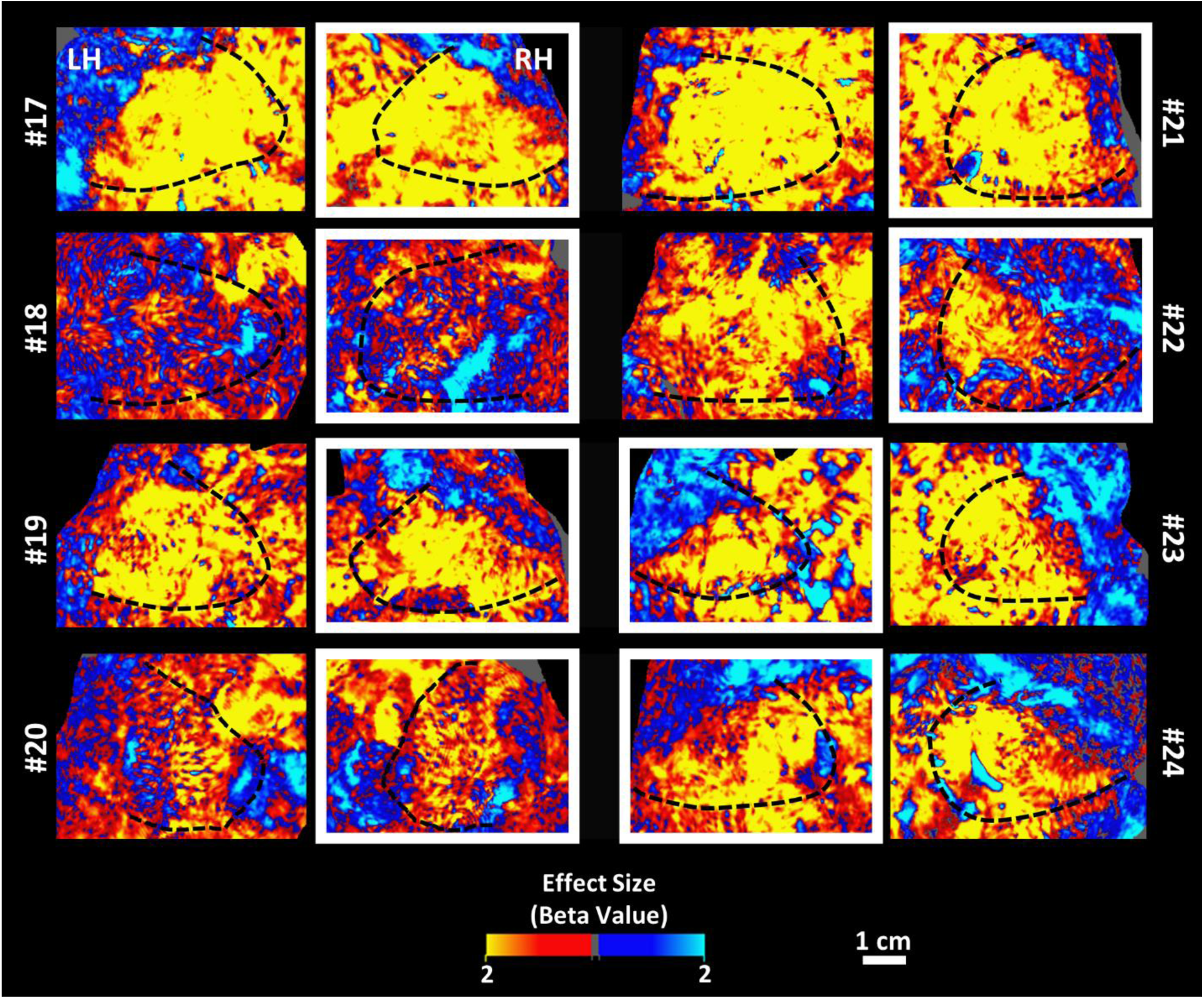
The OD activity mapping in 8 anisometropic participants, collected from the deep cortical depths. As in the strabismic participants (Fig. 4), the amplitude of OD response is larger relative to controls and the OD response extended beyond the V1-V2 borders. There was an overrepresentation of the fellow eye, as seen in strabismic participants. However, in contrast to the strabismic participants, this phenomenon was detected bilaterally without any apparent difference between the two hemispheres. Other details are the same as in Figs. 2 and 3.

As demonstrated in Fig. 4, in most strabismic individuals, we found larger regions that responded preferentially to the fellow eye within the hemisphere contralateral relative to fellow eye. However, participant #12 was the exception to this trend. Representation of the two eyes appeared to be more balanced in the hemisphere ipsilateral relative to the fellow eye (further analysis in Section 2.5). The fused activity patch close to the dorsal portion of V1-V2 border was readily apparent in 4 strabismic individuals (Participants #10, #12, #13 and #16), as in controls.

Among the participants categorized as having strabismic amblyopia, participants #9 and #14 had only a small difference in interocular visual acuity at the time of testing (≤0.12 logMAR; Table 1). Both individuals had a history of strabismus surgery and patching in childhood. Despite the small interocular visual acuity difference, both individuals showed signs of diplopia on Worth 4-dot testing, with reduced stereoacuity in participant #9 but not in participant #14. In both cases, we found an elevated OD response in V1, especially in the hemisphere contralateral to the fellow eye, similar to the other strabismic individuals. This result suggests that imbalanced ocular dominance may persist despite recovery of monocular visual acuity in the amblyopic eye, consistent with behavioral evidence for impaired *dichoptic* amblyopic eye visual acuity despite resolved interocular visual acuity differences after patching treatment (Birch, 2013; Birch et al., 2022).

In anisometropic individuals (Fig. 5), OD activity bias in favor of the fellow eye was detectable bilaterally in almost all subjects. There was no apparent difference between the two hemispheres. Among the four anisometropic individuals who did not show monocular suppression (Table 1), participants #17 and #22 showed a strong bias in favor of the fellow eye, but in participants #18 and #20 this bias was comparatively weaker. In contrast to strabismic individuals and controls, the activity patch along the V1-V2 border was less apparent in anisometropic individuals, likely due to strong bias in favor of the fellow eye.

Notably, the individual with deprivational amblyopia (participant #21) showed strong OD activity bias in favor of the fellow eye in both hemispheres, as in anisometropic individuals, even though the (unilateral; left eye) cataract was removed when the participant was a child, and the stimuli were perceived with best correction. Here again, this activity bias propagated into downstream visual areas. Considering the similarity between this individual’s OD pattern and those of anisometropic participants, we included this subject in the anisometropic group in the following analyses.

### 2.4. Reproducibility of the OD response maps across sessions

To compare the reproducibility of these maps across the three groups, we measured the correlation between OD activity maps evoked during the first and second scan sessions. This measurement was conducted separately for the activity evoked within the deep and superficial cortical layers and for the ipsilateral and contralateral hemispheres relative to the fellow eye. As we have shown previously (Nasr et al., 2016), OD activity maps remained highly correlated across sessions (Fig. 6). Two-way repeated-measures ANOVA (hemisphere and cortical depth, with a group factor), did not yield an effect of group (F(2, 21)=0.42, *p*=0.66)) or an interaction between group and the other independent variables (*p*>0.14). The same result was found in areas V2-V4, suggesting that activity maps were reproducible to the same extent for the three groups across visual areas.

**Figure 6.**
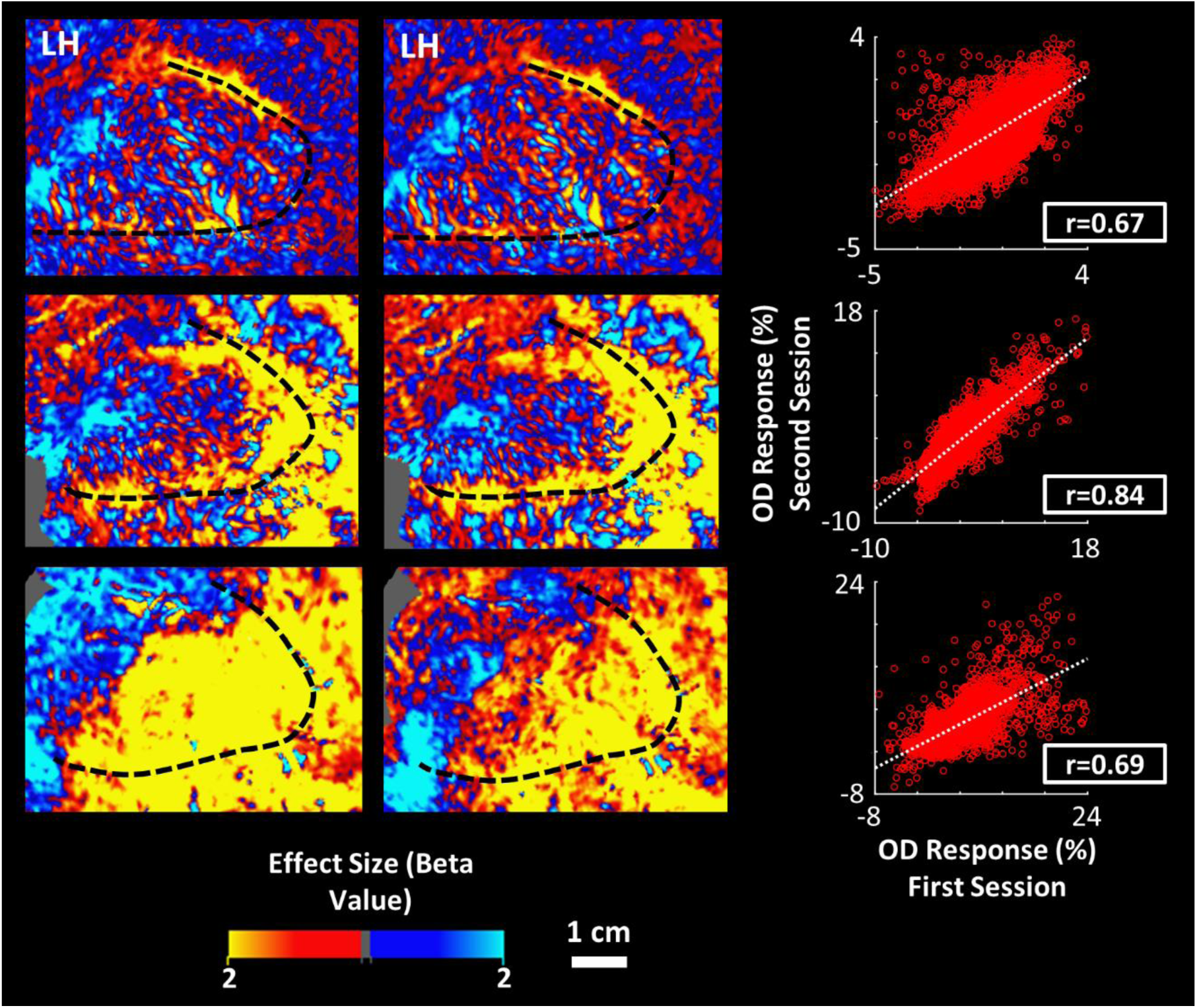
Reproducibility of the OD maps across scan session. The activity maps show the OD response evoked within the left hemisphere of the same control (top), strabismic (middle) and anisometropic (bottom) participants, as in Fig. 2, across two separate sessions (see Methods). The scatter plots highlight the correlation (*p*<10^-3^) between the OD response evoked with V1 across the two sessions. Each data point represents activity in one vertex from the reconstructed cortical surface mesh.

### 2.5. Overrepresentation of the fellow eye in amblyopic participants

Previous studies in human (Goodyear et al., 2002; Liu et al., 2004) and non-human primates (Smith III et al., 1997b; Kiorpes et al., 1998) have suggested an increased representation of the fellow eye in amblyopic compared to control participants. Consistent with these reports, we found an increase in the size of the V1 region that responded preferentially to the fellow eye in amblyopic participants compared to controls across deep and superficial cortical depth levels (Table 2). This effect also tended to be larger in anisometropic compared to strabismic individuals. Two-way repeated-measures ANOVA (hemisphere and cortical depth, with a group factor) yielded a significant effect of group (F(2, 21)=5.74, *p*=0.01), and a significant group × hemisphere interaction (F(2, 21)=3.86; *p*=0.04), but no group × cortical depth interaction (F(2, 21)=0.64; *p*=0.53) on the size of the V1 portion that responded preferentially to the fellow eye. Post-hoc analyses showed that in strabismic participants, this effect was significantly larger in the hemisphere contralateral compared to ipsilateral relative to the fellow eye (*p*=0.03). We did not find such a difference in either anisometropic (*p*=0.35) or control (*p*=0.56) participants. Notably, all measurements were normalized relative to the size of V1 area that was stimulated.

**Table 2.** The size of V1 portion that responded preferentially to the fellow/dominant eye (mean ± S. D.).

|  | Deep Cortical Layers | Superficial Cortical Layers |
| --- | --- | --- |
| **Control** | 55.37% ± 15.38% | 56.23% ± 15.54% |
| **Strabismic** | 66.80% ± 11.95% | 67.31% ± 12.15% |
| **Anisometropic** | 79.30% ± 15.30% | 78.53% ± 11.83% |

\* All measurements were normalized relative to the size of V1

### 2.6. The impact of amblyopia on the amplitude of the OD response

In addition to the change in the size of V1 portion that responded preferentially to the fellow eye, there was an increase in the amplitude of the evoked OD response in amblyopic compared to control participants (Fig. 7). A two-way repeated-measures ANOVA (as above) revealed a significant effect of group (F(2, 21)=11.91, *p*<10^-3^). Post-hoc analysis further showed that the evoked OD response in V1 was significantly larger in strabismic (*p*<10^-3^) and anisometropic (*p*<10^-5^) participants compared to controls without a significant difference between strabismic and anisometropic participants (*p*=0.22). Thus, in line with previous studies in humans (Conner et al., 2007) and non-human primates (Crawford and Von Noorden, 1979; Crawford et al., 1996; Smith III et al., 1997b; Kiorpes et al., 1998; Bi et al., 2011) studies, amblyopia increased the amplitude of the OD response in human V1 in our fMRI data.

**Figure 7.**
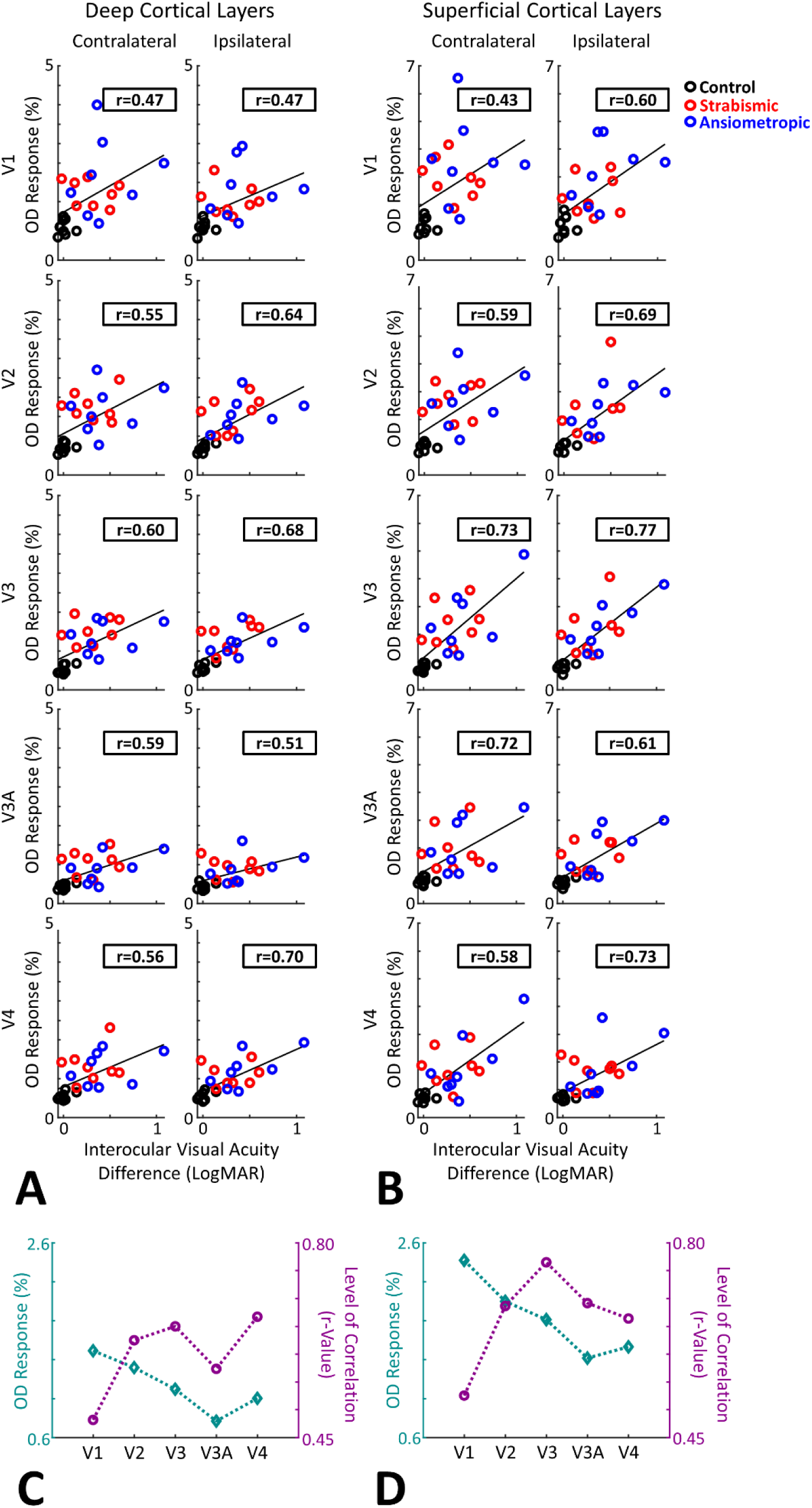
The amplitude of the OD response was measured in both deep (**A**) and superficial (**B**) cortical depths of V1-V4. Across all areas, the level of OD response was higher in the amblyopic participants compared to the controls, without a significant difference between the anisometropic and the strabismic individuals. To avoid signal cancelation, the ROI analysis was applied to the absolute value of OD response. Panels **C** and **D** show that, in both deep and superficial depths, the average OD response decreased in downstream visual areas relative to V1. However, the correlation between OD response and the interocular visual acuity difference increased from V1 to V2 to V3. Each point in these panels represents the average data from both hemispheres. Notably, the correlation values were calculated based on all participants. However, exclusion of controls did not change the overall results.

Importantly, the heightened OD response extended beyond V1 into downstream visual areas V2, V3, V3A and V4 (Fig. 7A and B). Despite a gradual decrease in the OD response amplitude from V1 through V4, the significantly stronger OD response in amblyopic individuals compared to controls was preserved across all tested areas (*p*<0.01). As in V1, the amplitude of the OD response remained comparable between strabismic vs. anisometropic participants (*p*>0.10). These results suggest that the impact of amblyopia on the OD response amplitude propagated to downstream visual areas, irrespective of amblyopia subtype.

Beyond this effect, we found a moderate correlation between the interocular visual acuity difference (as in the scans; see Table 1) and OD response amplitudes across visual areas V1-V4 (r>0.43; *p*<0.01). This correlation was considerably stronger in areas V2-V4 (r = 0.55 – 0.70) compared to V1 (r = 0.47), especially in deeper cortical depth levels, despite the decrease in the overall level of evoked OD response (Fig. 7C and D; Table 3). This correlation was similarly detected in contralateral and ipsilateral hemispheres and across superficial and deep cortical depth levels. To compare these correlation values more directly, we generated a linear multiple regression model using the interocular visual acuity difference as the dependent parameter and the OD activity evoked within V1-V4 (averaged between the two hemispheres) as the independent parameter. As demonstrated in Table 3, we found a stronger standardized beta value for V4 compared to V1 activity in both superficial and deep cortical depth levels. Thus, while the impact of amblyopia on the amplitude of the OD response was stronger in V1 compared to downstream visual areas, the correlation between OD response and interocular visual acuity difference was stronger in higher-level visual cortical areas such as V3 and V4.

**Table 3.** The correlation between the interocular visual acuity difference and OD activity evoked across V1-V4.

|  | Deep Layers |  |  |  |  | Superficial Layers |  |  |  |  |
| --- | --- | --- | --- | --- | --- | --- | --- | --- | --- | --- |
|  | V1 | V2 | V3 | V3A | V4 | V1 | V2 | V3 | V3A | V4 |
| **Correlation Value** | 0.48 | 0.62 | 0.65 | 0.57 | 0.67 | 0.53 | 0.69 | 0.77 | 0.69 | 0.66 |
| **Lower Confidence Interval** | 0.10 | 0.30 | 0.33 | 0.22 | 0.36 | 0.16 | 0.39 | 0.52 | 0.40 | 0.36 |
| **Upper Confidence Interval** | 0.74 | 0.82 | 0.83 | 0.79 | 0.84 | 0.77 | 0.85 | 0.89 | 0.86 | 0.84 |
| **Standardized Regression Coefficient** | 0.62 | 0.63 | 0.30 | 0.40 | 1.27 | 0.50 | 0.71 | 2.76 | 0.16 | 1.95 |

### 2.7. Contributions of residual strabismus

As reported in Table 1, the strabismic participants show some residual misalignments between the two eyes, despite prior surgical correction. To test whether mild strabismus, in the absence of amblyopia, may lead to the stronger OD responses we observed in individuals with strabismic amblyopia, we scanned a non-amblyopic individual with mild strabismus (separate from the other 8 controls (Participant #25; Table 1)). This participant showed normal, balanced visual acuities, no evidence of suppression or diplopia and showed measurable stereoacuity (70 seconds of arc).

As demonstrated in Fig. 8A, in this participant, the overall pattern of the OD response in V1 and downstream visual areas was distinguishable from that in individuals with strabismic amblyopia; instead, it more closely resembled the results in the controls (Fig. 2 and 3). Specifically, the size of the region that showed response preference to dominant eye stimulation within the contralateral (46.72%) and ipsilateral (43.03%) hemisphere (relative to the dominant eye) remained small compared to the individuals with strabismic amblyopia (Table 2). Similar results were detected within the superficial cortical levels. Thus, strabismus per se, in absence of amblyopia, is not the main cause of increased OD response in our participants. Notably, data from this participant were not used in any other analyses.

**Figure 8.**
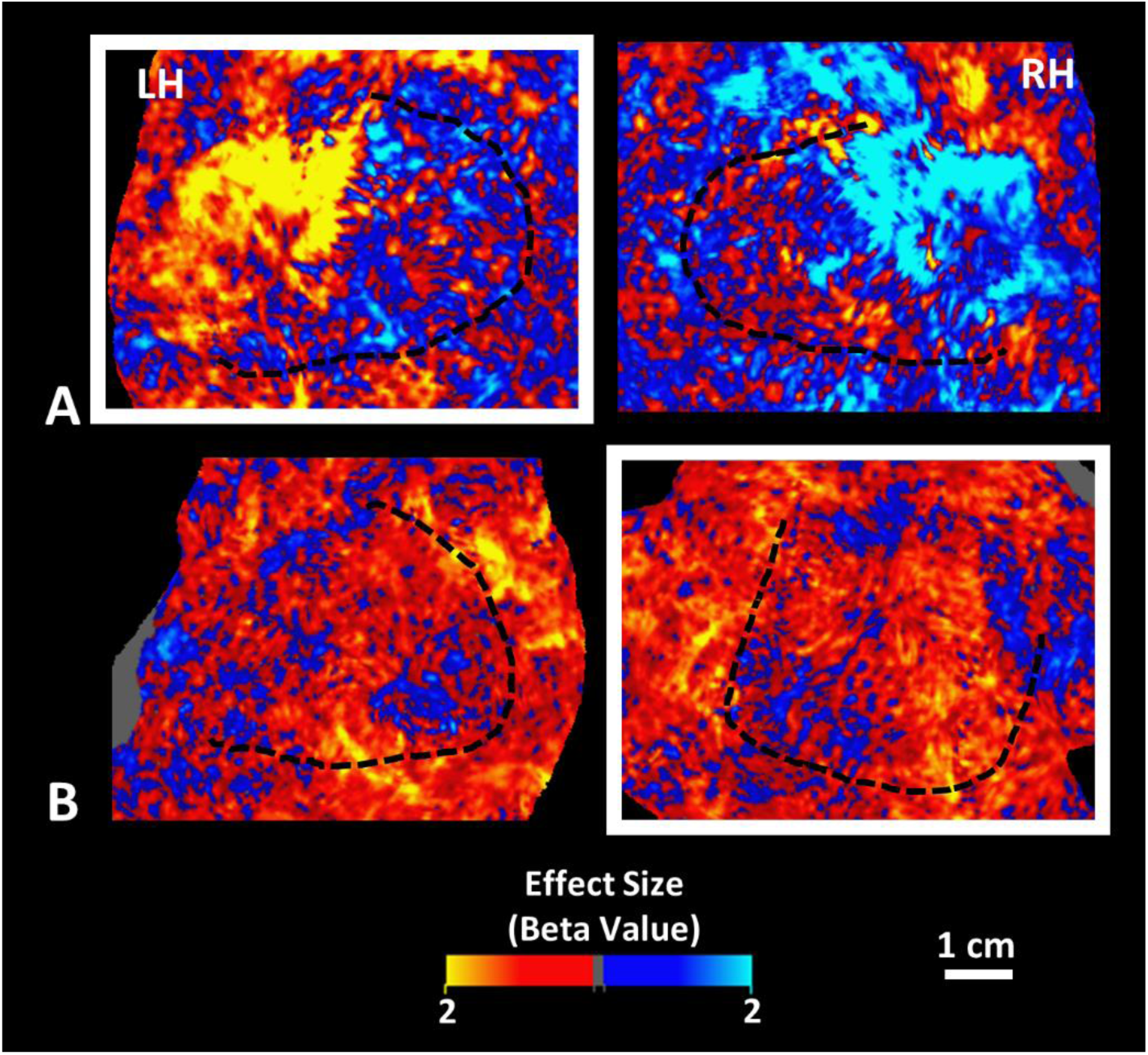
The OD activity mapping in non-amblyopic strabismic and anisometropic participants. Panel **A** shows the OD response in one non-amblyopic strabismic individual (participant #25; Table 1), collected from the deep cortical depth levels. The size of region that showed an OD bias in favor of the dominant eye remained close to what we found in control individuals (Table 2) in the contralateral (46.72%) and ipsilateral (43.03%) hemisphere (relative to the dominant eye). Panel **B** shows the OD response in one control subject (participant #6), after increasing the interocular visual acuity difference in favor of their dominant eye (from 0.06 to inducing 0.24 logMAR) by instructing the participant not to wear their contact lenses. Despite the increased level of interocular visual acuity difference, the evoked OD activity remained weaker compared to those detected in the amblyopic anisometropic individuals (Fig. 5). Other details are the same as Figs. 2-6.

### 2.8. Contributions of uncorrected visual acuity

Among the participants, two strabismic (Participants #12 and #15) and one anisometropic (Participant #22) individual could not be tested with their best optical correction. Even though this deviation had a relatively small impact on the level interocular visual acuity difference (Table 1; <0.11 logMAR), we tested whether this deviation from the best corrected visual acuity was the main source of increased OD activity in these individuals. In separate scan sessions, one control individual (Participant #6) was tested again with increased visual acuity difference by instructing the participant not to wear their prescribed contact lenses. Visual acuity worsened without correction by 0.76/1.00 (Right/Eye visual acuity) and the interocular visual acuity difference increased from 0.06 to 0.24 logMAR, as in the 3 participants with amblyopia who could not be tested with their best corrected visual acuity.

Fig. 8B shows the evoked OD response in this individual, measured within the deep cortical depth levels. While increased the level of bias in favor of the dominant eye, the level of evoked OD activity only increased 0.04% (fMRI signal level) and 0.12% in the contralateral and ipsilateral hemispheres, respectively. Moreover, OD activity in this participant was comparably weaker, compared to the OD activity detected on average in amblyopic individuals, in both contralateral (0.99%) and ipsilateral (0.82%) hemispheres. Similar results were detected in the superficial cortical depth levels. Thus, the increased OD activity in the three individuals with amblyopia who were unable to wear their best correction is only marginally attributable to the absence of optical correction.

### 2.9. The impact of amblyopia on the evoked response to binocular visual stimulation

In separate blocks, we also measured the evoked response to concurrent stimulation of both eyes. We compared binocular response amplitudes between regions preferring the dominant/fellow eye with those of the non-dominant/amblyopic eye for each group. As demonstrated in Fig. 9, results of this test revealed two important phenomena: First, there was no apparent difference between the level of response evoked within the V1 regions that responded preferentially to the fellow eye in amblyopic individuals compared to the V1 region that responded preferentially to the dominant eye in the controls. Second, in controls, binocular responses were comparable between V1 regions that responded preferentially to the dominant vs. non-dominant eye, whereas in amblyopic participants, evoked responses to binocular visual stimulation were stronger in V1 regions that responded preferentially to the fellow eye than those for the amblyopic eye. This effect appeared to be stronger at the superficial cortical depth, in the hemisphere contralateral to the dominant/fellow eye, and in anisometropic compared to strabismic individuals. Three-way repeated measures ANOVA (Hemisphere, Preferred-Eye, and Cortical Depth, with a group factor) yielded significant Group × Preferred-Eye (F(2, 20)=9.99, *p*<10^-3^), Group × Preferred-Eye × Layer (F(2, 20)=6.17, *p*<0.01), and Group × Preferred-Eye × Cortical Depth × Hemisphere (F(2, 20)=8.41, *p*<0.01) interactions for evoked binocular responses. A post hoc test to compare the response in anisometropic vs. strabismic individuals directly also showed a significant Group × Preferred-Eye × Cortical Depth (F(1, 14)=33.47, *p*<10^-3^) interaction whereas the other effects remained non-significant after correction for multiple comparisons. These results suggest that anisometropic, compared to strabismic, amblyopia is associated with a stronger decrease in the level of binocular activity within V1 regions that respond preferentially to the amblyopic eye.

**Figure 9.**
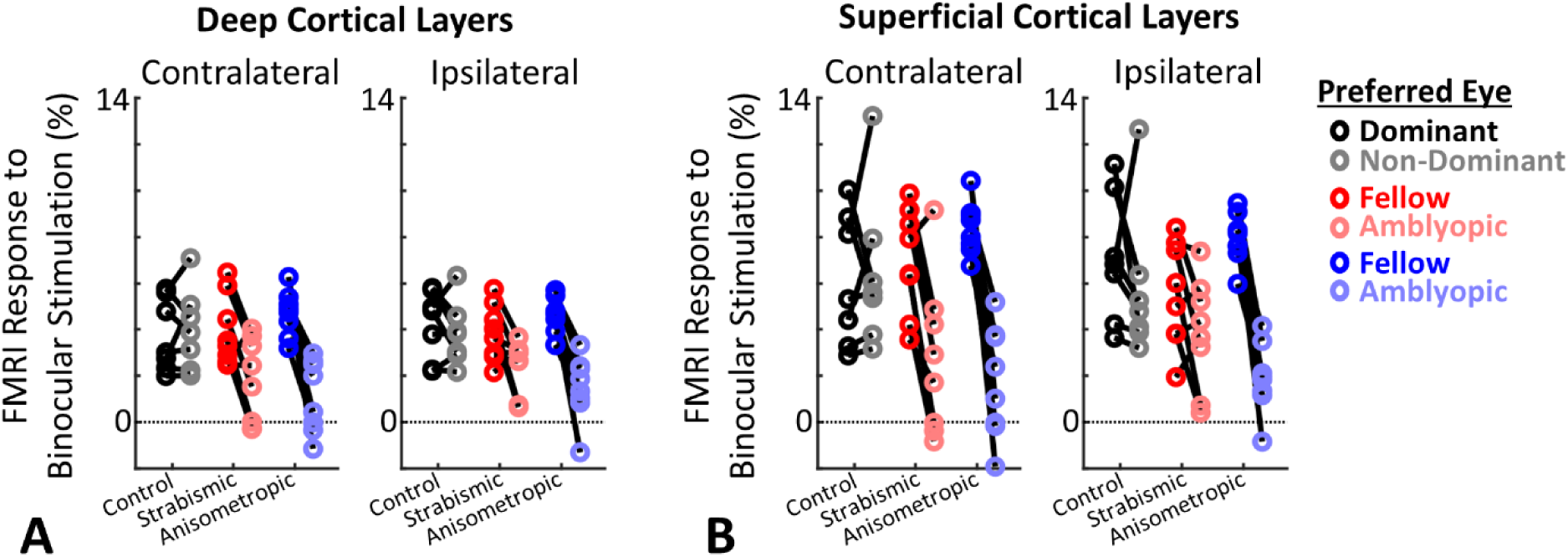
Activity evoked during binocular stimulation in V1 regions that responded preferentially to the dominant/fellow vs. non-dominant/amblyopic eye. Panels **A** and **B** show the activity evoked in deep and superficial cortical depth levels, respectively. In both depth levels and hemispheres, the level of activity evoked in V1 regions that responded preferentially to the dominant eye remained comparable across the three groups. Whereas, in V1 region that responded preferentially to non-dominant eye, binocular stimulation evoked a weaker response in anisometropic compared to strabismic and controls. This effect was more apparent in more superficial rather than deep cortical depths, and in contralateral rather than ipsilateral hemispheres (relative to the dominant eye). In all panels, each dot pair represents one individual subject.

### 2.10. The impact of amblyopia on the columnarity of OD response

Our knowledge of the impact of amblyopia on the vertical connections between cortical layers is limited to qualitative observations (Horton and Hocking, 1997). We tested the extent that amblyopia affects the functional link between deep and superficial cortical layers by comparing the correlation between activity maps evoked within deep and superficial cortical depths across the three groups.

As demonstrated in Fig. 10, we found an increased correlation between OD activity maps in deep and superficial cortical depth levels (i.e. inter-level correlation) of V1 in strabismic individuals compared to the two other groups. One-way repeated-measures ANOVA (hemisphere with a group factor) showed a significant effect of group (F(2, 21)=8.32, *p*<0.01) without any group × hemisphere interaction (F(2, 21)=0.29, *p*=0.75) for inter-level correlation values in V1. A post-hoc test showed that the magnitude of inter-level correlation was stronger in strabismic compared to anisometropic (*p*<0.01) and control (*p*<0.01) participants. Despite the extension of the OD response into the downstream visual areas (see above), application of this analysis to the evoked activity within V2-V4 did not yield a significant effect of group in any of those regions (*p*>0.17). This suggests that the impact of amblyopia on the columnarity of OD response is limited to primary visual cortex using these methods.

**Figure 10.**
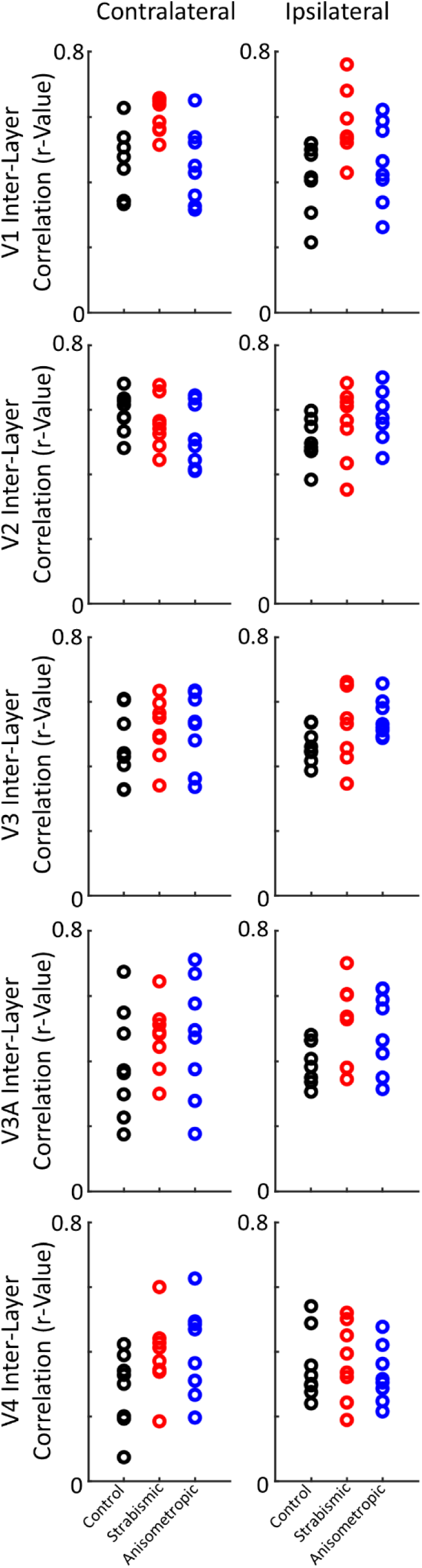
The level of correlation between the pattern of OD response evoked within deep and superficial cortical depths, across areas V1-V4. In area V1, but not the other visual areas, strabismic participants show a higher correlation compared to controls and anisometropic individuals. In each graph, each data point shows the data from one individual subject.

To test the reproducibility of this effect, first we repeated our tests for individual scan sessions, conducted on separate days, rather than the averaged activity maps. Again, two-way repeated measures ANOVA (hemisphere and session, with a group factor) showed a significant effect of group (F(2, 21)=7.74, *p*<0.01) without any significant group × session interaction (F(2, 21)=1.21, *p*=0.32) for inter-level correlation values, suggesting that the impact of amblyopia on the columnarity of the OD response was reproducible across scan sessions.

Second, in a randomly selected subset of our participants, consisting of 5 control and 4 strabismic individuals (Table 1), we tested whether this enhanced inter-level correlation is also seen in responses to dichoptically presented drifting gratings (rather than random dots). Briefly, we measured the level of OD activity evoked by gratings presented to fellow/dominant vs. amblyopic/non-dominant eye and then measured the correlation between the evoked OD activity across V1 at deep and superficial cortical depths. Despite fewer individuals participating in this test, we found significantly stronger inter-level correlation in strabismic compared to control participants (F(1, 7) = 11.09; *p*=0.01) and a significant group × hemisphere interaction (F(1, 7)=11.12, *p*=0.01). Thus, the enhanced inter-level correlation in the strabismic individuals was reproducible across stimulus types.

## 3. Discussion

By measuring the evoked activity in response to monocular and binocular stimuli, using high-resolution fMRI collected in an ultra-high field scanner, we showed direct evidence for the distinct impacts of anisometropia versus strabismus on the fMRI activity evoked within the human visual cortex. Specifically, we showed that the expanded representation of the fellow eye is more pronounced in anisometropic compared to strabismic participants, especially in the hemisphere ipsilateral relative to the fellow eye. Moreover, compared to strabismus, anisometropia has a stronger impact on the activity evoked during binocular stimulation within V1 regions that respond preferentially to the amblyopic eye. Strabismic amblyopia has a stronger impact on the level of correlation between the OD response evoked within V1 deep and superficial layers compared to anisometropic amblyopia. These findings were observed in anisometropic and strabismic participants with amblyopia of similar severity.

### 3.1. Consistency with findings based on animal models

Pronounced expansion of fellow eye representation in anisometropic compared to strabismic participants is consistent with single unit recordings in non-human primate V1 (Kiorpes et al., 1998; Bi et al., 2011). According to these studies, the number of neurons that respond preferentially to the fellow and amblyopic eye remains comparable in milder forms of strabismic amblyopia, whereas there is a relative increase of neurons responding preferentially to the fellow eye even in milder forms of anisometropic amblyopia.

The decreased binocular responses in our amblyopic participants is also consistent with previous reports that amblyopia may change the mechanism of interaction between the input from the two eyes (Smith III et al., 1997a; Kumagami et al., 2000; Bi et al., 2011; Farivar et al., 2011; Thompson et al., 2019). Here we showed that this decreased binocular activity is limited to V1 regions that respond preferentially to the amblyopic eye, at least by fMRI, suggesting that binocular integration is differentially impaired in V1 regions according to the ocular preference.

Our finding that strabismus is associated with an increase in the level of correlation between the OD activity in deep vs. superficial cortical depths is novel. To the best of our knowledge, no previous electrophysiological study had measured such a correlation in their participants directly. This finding is in line with anatomical studies in V1 of strabismic animals suggesting increased segregation between ODCs with opposite ocular preference (Shatz et al., 1977; Lowel, 1994; Tychsen et al., 2004). Moreover, according to animal studies, shrinkage of ODCs in layer 4 after monocular deprivation is associated with decreased cytochrome oxidase activity of blobs that fall in register with the shrunken columns, suggesting a change in vertical connections spanning cortical layers (Horton and Hocking, 1997). However, this effect has never been tested in vivo in anisometropic and/or strabismic participants.

### 3.2. Amblyopia impacts beyond V1

Since the original studies by Hubel and colleagues (Hubel et al., 1976), most amblyopia studies in animals have focused their efforts on understanding the impact of amblyopia on primary visual cortex. While this impact is expected to extend to downstream areas, only a few studies have examined this phenomenon in extrastriate visual cortex. Among them, Bi et al. reported that the increased OD response caused by strabismus extends to V2 (Bi et al., 2011). However, this extension was only detected in animals with severe amblyopia, suggesting a link between downstream extrastriate extension and visual impairment.

Consistent with that report, here we show that the correlation between the level of OD response and the interocular visual acuity difference, a functional measure correlated with ocular dominance shift, increased from V1 to downstream visual areas such as V3 and V4. This increase in correlation was detected despite the decrease in the OD response amplitude, suggesting that canonical propagation of OD deficits in amblyopia reflects functional visual impairment and highlighting the clinical relevance of downstream visual areas for future studies of evoked activity in the amblyopic brain.

### 3.3. Amblyopia impacts on visual attention

It could be argued that the reported correlation between the OD response and the interocular visual acuity difference is a result of amblyopia’s impact on the participant’s attention control mechanism, influencing both measurements concurrently. Degraded visual attention in amblyopia has been previously reported (Ho et al., 2006; Hou et al., 2016; Verghese et al., 2019). To reduce the influence of attentional bias that may confound OD responses and their correlation with visual acuity, two separate steps were taken: First, the OD response was measured while the participant’s attention was directed to an orthogonal task (i.e., shape change detection for the fixation object) separate from the stimuli used to elicit the OD measurement. Second, the fixation stimuli were presented dichoptically to reduce the potential impacts of biased attention in favor of the fellow eye. Thus, altered visual attention is unlikely to solely account for the strong OD response correlation with the visual acuity deficit in amblyopia.

### 3.4. The potential underlying mechanism for the increased OD response

Convergent evidence from both humans and non-human primates show that amblyopia is associated with an increase in the level of OD response in early visual areas. However, the mechanism underlying this phenomenon remains unclear. In several mammalian species, it is widely accepted that monocular deprivation in the first few weeks of life leads to a drastic decrease in afferent input originating from the amblyopic eye to V1 (Hubel et al., 1976; LeVay et al., 1980; Horton and Hocking, 1997). Given the typical developmental age of naturally occurring amblyopia (Shaw et al., 1988; Keech and Kutschke, 1995), disruption of binocular vision would not be expected to change the number of thalamocortical afferent inputs to V1. Consistent with those expectations, anatomical studies in humans (Horton and Stryker, 1993; Horton and Hocking, 1996) and animals (Horton et al., 1997) with naturally occurring amblyopia have reported intact ODC patterns in V1. Thus, the increased OD response among amblyopic human participants is unlikely to be attributable to decreased number of afferent connections originating from the amblyopic eye.

Amblyopia is also linked to changes in connections between ODCs. Anatomical studies have shown that strabismus and anisometropia are respectively associated with stronger and weaker segregation between ODCs (Shatz et al., 1977; Lowel, 1994; Tychsen et al., 2004). Horton and Hocking also showed evidence for a change in connections between layers 4 and 2/3 (the site of binocular convergence) after monocular deprivation (Horton and Hocking, 1997). However, the direction of this change (i.e. increased or decreased connectivity) remains unclear. Altered horizontal and/or radial connection between the ocular dominance columns may influence the ocular preference of V1 neurons and increase the OD response in amblyopic compared to non-amblyopic individuals. Longitudinal developmental studies are required to clarify the critical period for these effects and to test their correlation with the severity and distinct visual deficits of amblyopia.

### 3.5. Limitations

Despite recent advances in neuroimaging technologies (Polimeni et al., 2015; Blazejewska et al., 2019; Wang et al., 2022) that enabled us to map the OD response with relatively high spatial resolution (1 mm), our techniques may still have missed even-smaller OD patches, especially within the more peripheral portions of V1 (Adams et al., 2007). This caveat limits the interpretation of OD maps (Figs. 2-5). For instance, a relatively large patch that shows a uniform preference for one eye may contain small patches that are inaccessible due to limitations in spatial resolution.

Another limitation is that fMRI indirectly measures neuronal responses based on the concentration of deoxy-hemoglobin and blood flow. It has been shown that the existence of pial veins has a significant impact on increasing the level of evoked response and blurring the activity pattern in more superficial cortical layers (Koopmans et al., 2010; Polimeni et al., 2010; De Martino et al., 2013; Nasr et al., 2016). Existence of diving veins (Duvernoy et al., 1983) may also increase the level of correlation between deep and superficial depths. To the best of our knowledge, no previous study has shown evidence that amblyopia impacts vascularization of visual cortex. Nevertheless, the existence of diving veins may have influenced our estimation of the impact of amblyopia on the columnarity of OD response. Thus, any interaction between amblyopia and cortical depth must be assessed carefully and re-examined using fMRI sequences less sensitive to vascularization (Yacoub et al., 2007; Huber et al., 2015; Akbari et al., 2023). Unfortunately, these methods (e.g., spin echo and/or vascular space occupancy (VASO)) have low contrast-to-noise sensitivity that limits their application for assessing the mesoscale organization of the human brain.

Lastly, due to the small size of the head coil used in 7T scanners, we were not able to use accessories designed for lower field (e.g. 3T) scanners to correct visual acuity in those individuals who exclusively wore glasses, to stimulate more of the peripheral visual field (r<10°), and/or to monitor eye movements. While the impact of microsaccades and/or fixation instability on the fMRI signal is expected to be small, and our control experiments suggested that lack of optic correction and strabismus are unlikely to be responsible for the increased OD response in amblyopic subjects, these limitations prevented us from including individuals who required high degrees of optical correction and/or those who showed larger eye misalignments.

## 3.6. Conclusion

Despite its high prevalence in humans, our understanding of how amblyopia impacts the mesoscale organization of the visual system has been based primarily on animal models. In this study, high-resolution fMRI has documented the impact of amblyopia on the evoked OD response with functional correlates and drawn distinctions between the impact of anisometropia and strabismus on cortical responses.

## 4. Methods

### 4.1. Participants

Twenty-five human participants (10 females), aged 19–56 years old, participated in this study (Table 1). This included 7 anisometropic, one deprivational and 8 strabismic participants with amblyopia. We also included 8 individuals with normal (n=6) or correct-to-normal (n=2) visual acuity, as controls. One extra participant with mild strabismus (but no amblyopia) also participated in our study. The data from this individual is demonstrated separately. All participants had radiologically intact brains and no history of neuropsychological disorders.

During the main experiments, three amblyopic individuals could not wear their prescribed eye-glasses due to safety concerns with MRI compatibility (Table 1). To test the impact of this, one control participant underwent an additional control experiment during which the participant was tested without their contact lenses.

All experimental procedures conformed to NIH guidelines and were approved by Massachusetts General Hospital protocols. Written informed consent was obtained from all participants prior to all experiments.

### 4.2. Ophthalmological assessment

Outside the scanner, participants were tested by an optometrist (J.S.) with extensive experience with amblyopic individuals. During these tests, participants’ visual acuity (ETDRS retro luminant chart (Precision Vision)) was measured with pinhole (i.e. best corrected) and without pinhole (as in fMRI scans). The stereoacuity was measured using Randot stereo test (Stereo Optical). We identified the participant’s dominant eye (Miles Test) and tested for suppression or diplopia (Worth 4 Dot).

### 4.3. MRI experiments

Participants were scanned in an ultra-high field 7T scanner (whole-body system, Siemens Healthcare, Erlangen, Germany) for the functional experiments. All participants were also scanned in a 3T scanner (Tim Trio, Siemens Healthcare) for structural imaging.

During the fMRl experiments, stimuli were presented via an LCD projector (1024 × 768 pixel resolution, 60 Hz refresh rate) onto a rear-projection screen, viewed through a mirror mounted on the receive coil array. MATLAB 2021a (MathWorks, Natick, MA, USA) and the Psychophysics Toolbox (Brainard, 1997; Pelli, 1997) were used to control stimulus presentation. The participants were instructed to look at a centrally presented fixation object (radius = 0.15°) and to do either a shape-change for the fixation target (circle-to-square or vice versa) during the OD measurements or a random dot-detection during the retinotopic mapping. These tasks were conducted without any significant difference across experimental conditions (*p*>0.10).

#### 4.3.1. Response to monocular and binocular visual stimulation based on moving random dots

All participants completed 2 separate scan sessions. In each session, we stimulated the participant’s fellow (dominant) and amblyopic (non-dominant) eyes in different blocks (i.e., block-design; 24 s per block). The stimuli were sparse (5%) moving random red (50% of blocks) and green (the rest of blocks) dots (0.09° × 0.09°; 56 cd/m^2^), presented against a black background. In separate blocks, we also measured the response to binocular presentation of the simultaneous stimulation of both eyes (with zero disparity) in all participants except for one control.

Participants viewed the stimuli through custom made anaglyph spectacles (with red and green filters) mounted to the head coil. During the blocks, dots were oscillating horizontally (-0.22° to 0.22°; 0.3 Hz). Stimuli extended 20° × 26° in the visual field. Each experimental run began and ended with 12 s of uniform black. The sequence of blocks was pseudo-randomized across runs (14 blocks per run) and each participant participated in 12 runs. Filter laterality (i.e., red-left vs. red-right) was counter-balanced between sessions and across participants.

#### 4.3.2. Response to monocular visual stimulation based on moving gratings

To test whether the strabismic amblyopia impact on the columnarity of the OD response was detectable based on stimuli other than random dots, in this experiment participants were presented with gratings (2.25 cycle/degree). Red and green gratings were presented in different blocks (24 s per block) and participants viewed the stimuli through custom anaglyph spectacles mounted on the head coil. To avoid adaptation, gratings were oscillating left-to-right (-0.22° to 0.22° (0.3 Hz)). Stimuli were presented against a black background, extending 20° x 26° in the visual field. The orientation of gratings varied randomly between blocks.

Each experimental run began and ended with 12 s of uniform black. The sequence of blocks was pseudo-randomized across runs (7 blocks per run) and each participant participated in 2 runs. Filter laterality (i.e., red-left vs. red-right) was counter-balanced across participants.

#### 4.3.3. Retinotopic mapping

For all participants the border of retinotopic areas were defined retinotopically (Sereno et al., 1995). Stimuli were based on a flashing radial checkerboard, presented within retinotopically limited apertures, against a gray background. These retinotopic apertures included wedge-shaped apertures radially centered along the horizontal and vertical meridians (polar angle = 30°). These stimuli were presented to participants in different blocks (24 s per block). The sequence of blocks was pseudo-randomized across runs (8 blocks per run) and each participant participated in at least 4 runs.

### 4.4. Imaging

Functional experiments (see above) were conducted in a 7T Siemens whole-body scanner (Siemens Healthcare, Erlangen, Germany) equipped with SC72 body gradients (70 mT/m maximum gradient strength and 200 T/m/s maximum slew rate) using a custom-built 32-channel helmet receive coil array and a birdcage volume transmit coil. Voxel dimensions were nominally 1.0 mm. We used single-shot gradient-echo EPI to acquire functional images with the following protocol parameter values: TR=3000 ms, TE=28 ms, flip angle=78°, matrix=192×192, BW=1184 Hz/pix, echo-spacing=1 ms, 7/8 phase partial Fourier, FOV=192×192 mm, 44 oblique-coronal slices, acceleration factor *R*=4 with GRAPPA reconstruction and FLEET-ACS data (Polimeni et al., 2015) with 10° flip angle. The field of view included occipital cortical areas V1, V2, V3 and the posterior parts of V4v and V4d.

Structural (anatomical) data were acquired in a 3T Siemens TimTrio whole-body scanner, with the standard vendor-supplied 32-channel head coil array, using a 3D T1-weighted MPRAGE sequence with protocol parameter values: TR=2530 ms, TE=3.39 ms, TI=1100 ms, flip angle=7°, BW=200 Hz/pix, echo spacing=8.2 ms, voxel size=1.0 × 1.0 × 1.33 mm^3^, FOV=256 × 256 × 170 mm^3^.

### 4.5. General data analysis

Functional and anatomical MRI data were pre-processed and analyzed using FreeSurfer and FS-FAST (version 7.11; http://surfer.nmr.mgh.harvard.edu/) (Fischl, 2012).

#### 4.5.1. Structural analysis

For each participant, inflated and flattened cortical surfaces were reconstructed based on the high-resolution anatomical data (Dale et al., 1999; Fischl et al., 1999; Fischl et al., 2002). Then, during this reconstruction process, the standard pial surface was generated as the gray matter border with the surrounding cerebrospinal fluid or CSF (i.e. GM-CSF interface). The white matter surface was also generated as the interface between white and gray matter (i.e., WM-GM interface). To enable intra-cortical smoothing (see below), we also generated a family of 9 intermediated equidistant surfaces, spaced at intervals of 10% of the cortical thickness, between WM-GM and the GM-CSF interface surfaces. To improve the co-registration of functional and structural scans, all surfaces were unsampled (Wang et al., 2022).

#### 4.5.2. Functional analysis

The collected functional data were first unsampled (to 0.5 mm isotropic) and then corrected for motion artifacts. For each participant, functional data from each run were rigidly aligned (6 DOF) relative to their own structural scan using rigid Boundary-Based Registration (Greve and Fischl, 2009). This procedure enabled us to compare data collected for each participant across multiple scan sessions.

To retain the spatial resolution, no tangential spatial smoothing was applied to the imaging data acquired at 7T (i.e., 0 mm FWHM). Rather we used the more advanced method of radial (intracortical) smoothing (Blazejewska et al., 2019) – i.e., perpendicular to the cortex and within the cortical columns. For deep cortical depths, the extent of this radial smoothing was limited to WM-GM interface and the adjacent 2 surfaces right above it (see above) – i.e., the bottom 30% of the gray-matter thickness starting from the WM-GM interface. For the superficial cortical depths, the extent of this procedure was limited to GM-CSF interface and the adjacent 2 surfaces right below it. For the middle cortical layers, used only for presentation (Fig. 2), the extent of this procedure was limited to the three middle reconstructed cortical surfaces.

A standard hemodynamic model based on a gamma function was fitted to the fMRI signal to estimate the amplitude of the BOLD response. For each individual participant, the average BOLD response maps were calculated for each condition (Friston et al., 1999). Finally, voxel-wise statistical tests were conducted by computing contrasts based on a univariate general linear model, and the resultant significance maps were projected onto the participant’s anatomical volumes and reconstructed cortical surfaces.

### 4.6. Region of interest (ROI) analysis

To test the impacts of amblyopia on the OD response, ROIs including deep and superficial depths of areas V1, V2, V3, V3A, and V4, defined for each participant based on their own structural and retinotopic mapping (see above).

To test the impact of amblyopia on the evoked response to binocular stimulation, V1 surface was divided into two ROIs based on the ocular preference of the vertices, defined during the monocular tests. These ROIs were defined independently for deep and superficial cortical depths.

Notably, no hemisphere was excluded from any ROI analyses and all vertices within each ROI were used in the analyses.

### 4.7. Statistical data analysis

Three independent parameters included group (anisometropic vs. strabismic vs. control participants), hemisphere (ipsilateral vs. contralateral relative to the dominant/fellow eye) and cortical depth level (deep vs. superficial). To test the impact of these parameters, we used either one-way or two-way repeated-measures ANOVA with a group factor. Since this analysis is particularly susceptible to the violation of sphericity assumption, caused by the correlation between measured values, when necessary (determined using a Mauchly test), results were corrected for violation of the sphericity assumption, using the Greenhouse-Geisser method. All post-hoc analyses were conducted after Bonferroni correction for multiple comparisons.

### 4.8. Data availability statement

Data and codes will be shared upon request.

## Acknowledgment

This work was supported by NIH NEI (R01 EY030434, EY029713 and K08 EY030164), and by the MGH/HST Athinoula A. Martinos Center for Biomedical Imaging. Crucial resources were made available by a NIH Shared Instrumentation Grant S10-RR019371. We thank Dr. Jason Stockmann and Ms. Azma Mareyam for helping with hardware maintenance during this study. We thank Ms. Amanda Nabasaliza for her help with the recuitment. We also thank Drs. Daniel Tso and Jonathan Horton for their helpful comments.

